# Human cerebral tissue growth is a critical process

**DOI:** 10.1101/2022.04.28.489779

**Authors:** Egor I. Kiselev, Florian Pflug, Arndt von Haeseler

## Abstract

We develop a Fokker-Planck theory of tissue growth with three types of cells (symmetrically dividing, asymmetrically dividing and non-dividing) as main agents to study the growth dynamics of human cerebral organoids. Fitting the theory to lineage tracing data obtained in next generation sequencing experiments, we show that the growth of cerebral organoids is a self organized critical (SOC) process. We derive analytical expressions describing the evolution of clonal lineage sizes and discuss possible organizational mechanisms behind the critical growth.

## I. INTRODUCTION

The mechanisms of tissue growth and renewal are a core topic of stem-cell biology [1, 2]. In particular, the role of stochasticity in cell differentiation is discussed [3–8]. Modern gene sequencing protocols – commonly referred to as next generation sequencing (NGS) – allow to study the genome of individual cells (single cell sequencing) [9]. In combination with the labeling of cells with inheritable DNA sequences, these techniques enable large scale, quantitative studies of cell populations in biological tissues, allowing to trace back offspring populations to their individual ancestral cells [10, 11]. Such lineage tracing experiments have revealed that offspring numbers in mammalian cerebral tissue can vary by several orders of magnitude [6, 12], which supports the hypothesis that stochasticity is an important property of cell proliferation and differentiation in the developing cerebral cortex.

Cerebral organoids are models of the human cerebral cortex grown from stem cells, and are used to study neural development and diseases [13, 14]. This work presents a study of lineage tracing data obtained by sequencing 15 organoids at different stages of their development [12]. We take a physics point of view on the population dynamics of cell lineages, and show that organoid growth is a self organized critical (SOC) process. Self organized criticality [15, 16] is the working principle behind a multitude of phenomena in physics [17, 18], geology [19–22], biology [23–27] and the social sciences [28–30]. One hallmark of SOC and criticality in general is the appearance of power-law distributions. A prime example is the Gutenberg-Richter-law in seismology stating that the number of earthquakes having an energy *E* is distributed according to *N* (*E*) ≈ *E*^−1−*b*^ with *b* ≈ 0.5 [20, 31] (see also footnote 2 in Ref. [20] and Ref. [32] for alternative formulations of the law). The number of neurons *n* participating in a neural avalanche is known to scale according to the power-law *n^−τ^* with *τ* ≈ 1.5[24, 33]. Other prominent examples from biology are the near critical state of hair cells in the inner ear [34, 35] and the collective dynamics of bird flocks and insect swarms [36–38].

SOC was first introduced using the sand pile as a paradigmatic example [15]: Imagine a table onto which sand grains are dropped at a constant rate. A sand pile begins to grow and its slope slowly increases. The growth continues until the pile occupies the whole table and its slope reaches a critical value. At this point, the rate at which grains fall off the table is equal to the rate at which they are dropped onto the pile. This is the critical state: if the slope of the sand pile is further increased, the sand pile will become unstable and collapse. For our purposes, the self organized branching process proposed in Ref. [40] is a good means to describe the sand pile at this stage. It provides a simple picture of how the sand pile organizes itself towards the critical state: when a new grain is dropped onto the pile, it can either (a) trigger another grain to fall downhill, or (b) stick to the surface. Let the probabilities for the two outcomes be *p* and 1 – *p*, respectively. If (a), the triggered grain can continue the process by triggering other grains in a chain reaction or *avalanche.* This model is illustrated in Fig. 1. Each grain possesses a typical potential energy that is set free whenever event (a) takes place. However, when grains fall off the table, potential energy leaves the system. On average, each grain in the pile will now have less potential energy, and the probability *p*, that it can be triggered decreases. However, if *p* is low and no grains leave the table, the overall potential energy of the pile increases, and *s*_0_ does *p* in the next iteration of the process. It is shown in Ref. [40] that the system will approach a state with *p* ≈ 1 – *p* ≈ 1/2- the critical state. In particular, the avalanche size, i.e. the number *n* of grains leaving the table in each iteration of the process, will be distributed according to *n*^−3/2^. Experiments with rice piles demonstrated that avalanche sizes are indeed distributed according to power-laws, albeit with somewhat different exponents [39]. This is due to the fact, that the simple branching model neglects multiple interactions between grains. However, the appearance of the 3/2 exponent makes the critical branching process an attractive model for the description of neural avalanches [33] and earthquake propagation [21, 41, 42].

**Figure 1.**
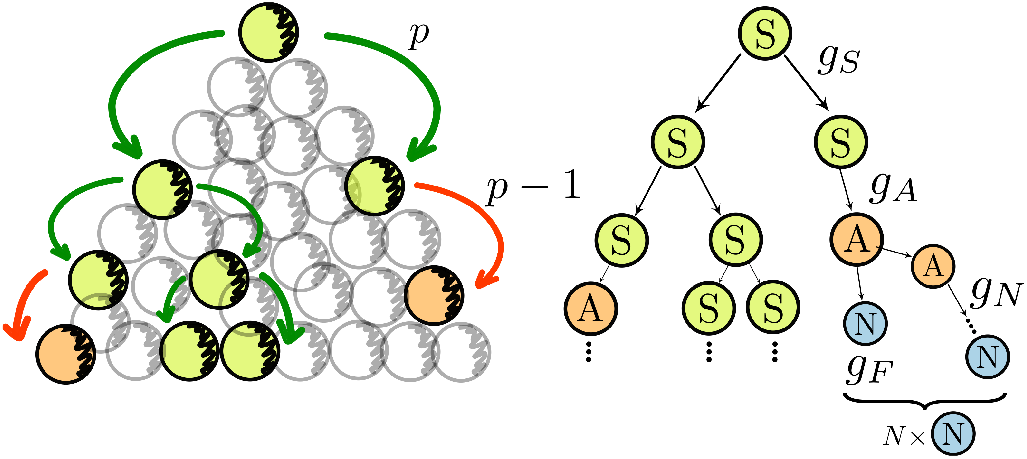
Avalanches in self organized critical (SOC) processes and stem cell division and differentiation can be mapped onto each other. The left part shows a schematic representation of an avalanche propagating down a SOC system, e.g. a sand- or rice-pile [15, 16, 39]. Green grains indicate active sites carrying the avalanche downhill, while orange grains are dead-ends which remain inactive. Criticality is reached when the probability of triggering an active site and reaching a dead end are equal: *p* = 1 – *p* = 1/2 [40]. The right part shows a system of dividing and differentiating cells. While stem cells (green, S) maintain the potential for symmetric division (S→2S) and are analogous to the active sites of the right hand SOC process, differentiated cells (orange and blue) either divide asymmetrically (A→A+N) or loose this ability altogether (N). In our model, S-cells drive the SOC avalanche while each asymmetrically dividing cell produces *N* N-cells on average, until it is terminated by a direct A→N process. At criticality, the rates *g_S_* (division) and *g_A_* (differentiation) must be equal.

Our study begins with the observation that the numbers of descendants of an individual stem cell in the organoid (lineage sizes) are roughly distributed according to a 3/2-power-law (Fig. 2). This behavior becomes more and more pronounced at late stages of the organoid development. However, to study tissue growth, a more sophisticated, dynamical branching model than the one described above is required. The three agents of our model, which we call the *SAN model,* are symmetrically dividing S-cells that can be considered as stem cells, asymmetrically dividing A-cells and non-dividing N-cells. S-cells undergo symmetric division (S→2S), differentiation (S→A) and death (S→0). A-cells have committed to a developmental trajectory and produce N-cells through asymmetric divisions (A→A+N) until the process is terminated by direct differentiation (A→N) or death. The branching process of symmetric division and differentiation of S-cells with rates *g_S_* and *g_A_* is at the heart of the SAN-model. Criticality is reached, when the two rates are equal: *g_S_* = *g_A_*. Despite being a sufficiently crude representation of biological reality, in an extensive computational experiment, the SAN-model was found to describe organoid growth at later stages very well[43]. The model and its connection to the SOC branching model is illustrated in Fig. 1. We solve the SAN-model analytically in the continuum limit and show, in different ways, how 3/2-power-law distributions of cell populations arise near criticality. Fitting the model predictions to the empirical data of Ref. [12], we show that organoid growth is indeed a self organized critical process (see Fig. 3 and Sec. II C). A crucial aspect of our theory is that it describes the SOC of organoid growth as a dynamical process in order to capture organoids at different developmental stages. This is in contrast to many other mean field SOC models, e.g. Ref. [40].

**Figure 2.**
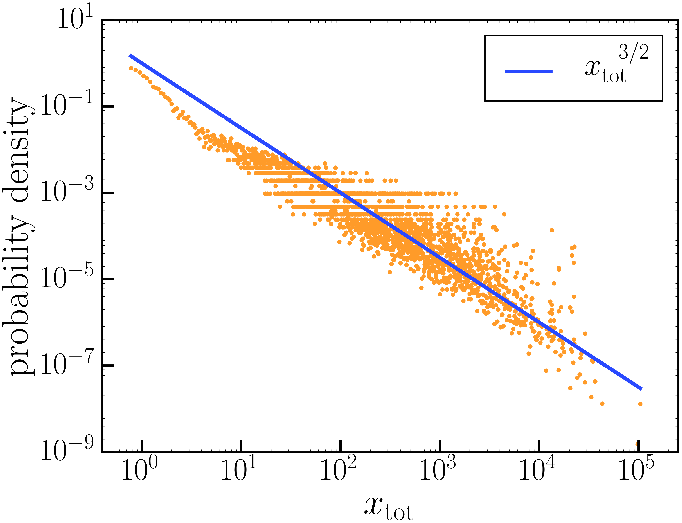
Lineage size xtot distribution of a single organoid at day 40 (see Sec. II C for details). The distribution roughly follows a critical 3/2-power-law.

**Figure 3.**
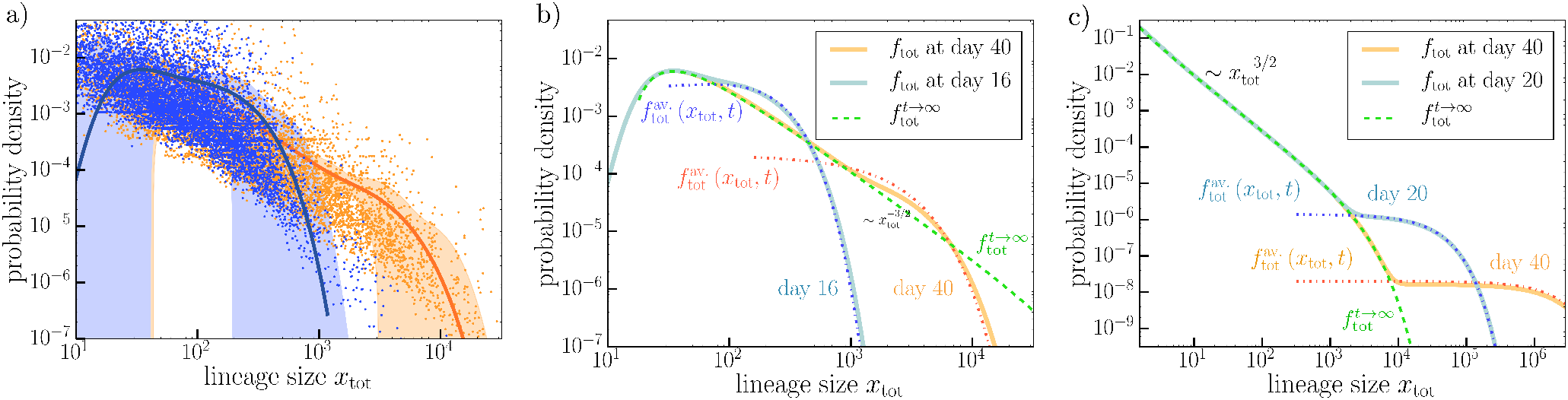
a) lineage size probability density of the SAN model *f*_tot_ (*x*_tot_, *t*) at *t* = 16 days and *t* = 40 days (solid lines) and the empirical probability densities at the same days (scattered points). The shown data is combined from six sequenced organoids (three per day). The estimates for the SAN parameters *α* (net growth rate) and *β* (stochasticity rate) are given in Table I. *f*_tot_ (*x*_tot_, *t*) was obtained using a Fast Fourier Transform (FFT) of the characteristic function of Supplementary Eq. (S 33). b) FFT-calculated SAN model probability densities form a) and the analytical approximations 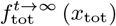 (dashed, green line) and 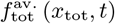 of Eqs. (15), (16) for the parameter estimates of Table I (*α* < 0). For *t* → ∞ the distribution approaches the weakly truncated 3/2-power-law Levy distribution 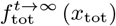 everywhere. For small *t* there is a region where *f*_tot_ (*x*_tot_, *t*) is well approximated by 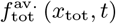 and exceeds 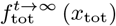 before it quickly truncates. This region moves towards higher *x*_tot_ as *t* increases and marks the active part (avalanche part) of the population with actively deviding S-cells. Since the net S-cell growth rate *α* is negative, this region becomes less and less pronounced for large *t*. c) SAN model predictions for small *α* > 0. The S-cell population proliferates indefinetly, however *f*_tot_ (*x*_tot_, *t*) still approaches 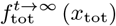, except for very large lineage sizes. Here the probability distribution exceeds the *t* → ∞ limit. This region marks the avalanche regime of active S-cell proliferation for positive *α*. We used *α* = 0.2, *β* = 10, *s*_0_ = 1 and *N* =1.

In Section II, we present the results of this paper. Sections II A and II B explain, in a nontechnical manner, the implications of SOC tissue growths and the origins of power-law lineage size distributions. Section II C presents the experimental results which show that human cerebral tissue growth is indeed a SOC process. In Sections II D and II E we summarize the main mathematical results: the SAN-model and its analytical solutions which shade light on the growth dynamics and the qualitative features of lineage size distributions for different values of the model parameters. In Section II F a more intuitive picture of lineage dynamics based on path integrals is presented. This approach explains why, in the SOC scenario, the evolution of large lineages is essentially a universal process which does not depend on initial conditions. Finally we discuss our results and possible sources of error in Sec. III.

## II. RESULTS

### A. What critical tissue growth implies

Several lessons can be learned from applying the branching paradigm to the problem of cerebral tissue growth: First, SOC implies the presence of a mechanism that organizes and maintains the critical state. In it’s original formulation in Ref. [40], the SOC branching process is a highly nonlocal phenomenon. Returning to the sand pile example for a moment, the behavior of each grain depends on the critical state of the whole sand pile. Similarly the behavior of each S-cell in the organoid must be regulated such that the overall balance between the rate *g_S_* of divisions and *g_A_* of differentiations is maintained and the organoid is kept at criticality. In the sand pile model, this balance is maintained by the loss and gain of total potential energy in the system. In organoids a similar mechanism could be at work, with potential energy replaced by competition for a different quantity in limited supply, such as space, nutrients or a chemical signal. The balancing mechanism does not necessarily have to be global – local rules which that involve only single active sites and their immediate descendants have been shown to be sufficient for maintaining criticality [42, 44, 45]. However, whether global or local, the mechanism likely involves an extracellular component or property. Otherwise, the balance between divisions and differentiations would have to be maintained through an innate propensity of stem cells to differentiate spontaneously, which contradicts stem cells being capable of indefinite self renewal [46].

Second, tuning itself to the critical point the organoid maximizes the stochasticity of the growth process. This is a striking observation: we are not simply dealing with stochastic growth at a given rate. Instead, the growth is fully determined by stochasticity and there is no fundamental characteristic time scale controlling the process. This is in contrast to many other examples of organ growth and regeneration [47–49] were the growth process is arrested in a coordinated manner. An important feature of critical growth is that the long term dynamics of large lineages, which make up most of the organoid, does not depend on the initial conditions of the process. Here stochastic fluctuations have erased all memory of the lineage’s initial size. This is illustrated if Fig. 4 and we dwell on this point in Sec. II F. This is possibly an important advantage of the critical regime: the outcome of the growth process is less influenced by perturbations in its initial stages, when the tissue is most susceptible to disturbances.

**Figure 4.**
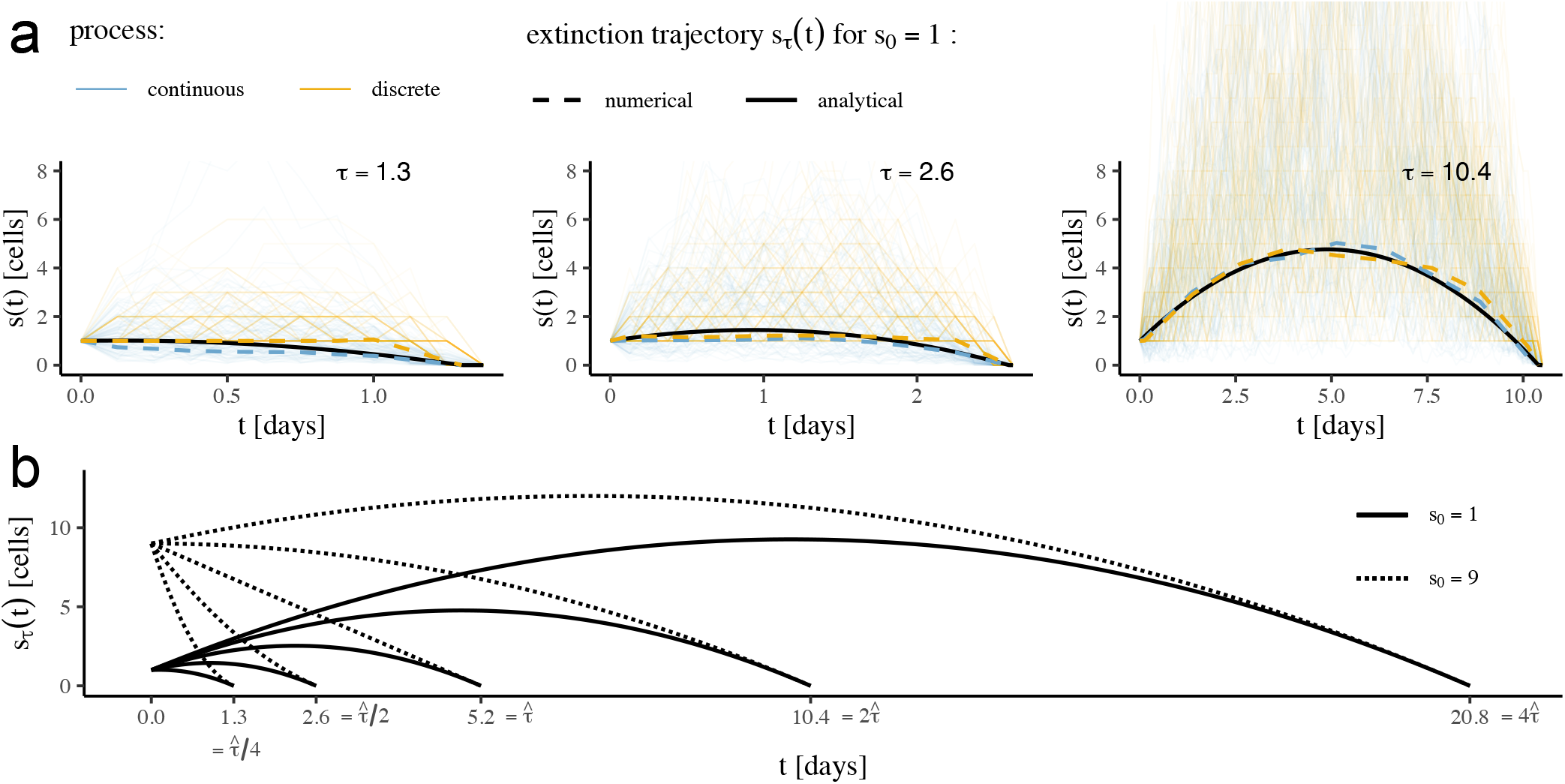
a) Theoretical (solid line) extinction trajectory *s_τ_* (*t*) and simulated (dashed lines) extinction trajectories for the continuous (blue) and discrete (yellow) process for *τ* = 1.3 *τ* = 2.6, and *τ* = 10.4 for parameters *s*_0_ = 1, *g_S_* = *g_D_* = 2 (i.e. *β* = 4, 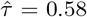). Simulated trajectories were computed by selecting the 10,000 trajectories with extinction time closest to *τ* from 2 · 10^6^ simulated trajectories of the respective process, evaluating these trajectories at 9 equally spaced times to obtain 10,000 samples of a 9-dimensional multivariate distribution, and locating the mode using the mean-shift algorithm, b) Theoretical extinction trajectories for initial population sizes *s*_0_ = 1 (solid lines) and *s*_0_ = 9 (dotted lines) and different extinction times *τ* (again *β* = 4). For *s*_0_ = 9, the critical time is 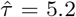, and the different trajectories exhibit the *sublinear, superlinear* and *initial growth* cases explained in the main text. The larger the extinction time *τ*, the more do the trajectories for *s*_0_ = 1 and *s*_0_ = 9 approach one another for large *t*. This demonstrates, that in the critical state the trajectories are dominated by stochasticity, and hence obtain a universal form independent of the initial conditions.

### B. How birth-death dynamics gives rise to power-laws

To show how the process of stem cell proliferation and differentiation can organize itself towards criticality and produce power-law distributions, let us first study a simplified model. In this model we consider stem cells (S-cells) which symmetrically divide with a rate *g_S_*, and differentiate at a rate *g_D_*. Once differentiated, the cells do not proliferate any more in this simple model. Since at each moment in time the cells can either evolve, divide, or simply do nothing, the process is a stochastic. The stem sell population is therefore described by a probability density function *f* (*s*), where *s* is the number of S-cells. The dynamics of the distribution function *f* (*s*) is described by the well known birth-death Fokker-Planck equation [50, 51]:

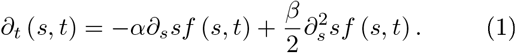

Here *s* is treated as a continuous cell count and the constants are defined as *α* = *g_S_* – *g_D_* and *β* = *g_S_* + *g_D_*.

Regarding the treatment of cell counts as a continuous quantity, we remark that if *s* is sufficiently large for a change by one to be considered infinitesimal compared to s, the continuous process specified by the Fokker-Planck equation 1 and the corresponding discrete process will be equivalent. For small *s* the laws will differ, yet we emphasize that both are reasonable approximations of biological reality. While using a discrete version of the process may seem preferential at first because it ensures integral cell counts, it does *s*_0_ at the cost of treating cells as memoryless and cell division as spontaneous. But in reality, cells are not memoryless, and cell division is only the end result of a series of changes to a cell’s internal state as it transitions through the G1, S, G2 and M stages of the cell cycle. Because the continuous process accounts for these changes by allowing cell counts to change gradually, it may be in fact the model that is closer to reality.

Eq. (1) was introduced by Feller [52] to study the dynamics of self replicating populations. The first right hand side term is a drift term that increases (*α* > 0) or diminishes (*α* < 0) the population sizes, while the second term is responsible for the broadening of the probability density due to stochastic processes, *α* can be thought of as a net growth rate. Since differentiation events diminish the S-cell population *g_D_* enters *α* with a negative sign. Both differentiation and symmetric division contribute to the system’s stochasticity. Hence, both enter *β* with a positive sign. Solving Eq. (1) with the initial condition *f* (*s,t* = 0) = *δ* (*s* – *s*_0_), which corresponds to *s*_0_ S-cells at *t* = 0, the expected S-cell population evolves according to [52]

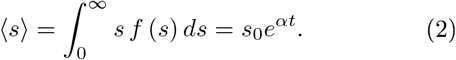

For *α* > 0, the population is exponentially growing. For *α* < 0 it is shrinking and will certainly die out. The critical case is *α* = 0. Interestingly at *α* = 0, even though the expected population size is constant, a single population also certainly dies out eventually (or, in our terms, differentiates). The probability that a population has died out at time *t* is then given by [52]

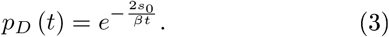

In the following, we demonstrate that tissue growth in cerebral organoids does organize itself towards criticality, and that therefore the rates of differentiation and proliferation have to be autonomously fine-tuned to ensure that *α* = *g_S_* – *g*_0_ = 0. A crucial sign of critical behavior is the appearance of power-laws [15–17]. For cerebral organoids, we find experimentally[12, 43] that the distribution *f* of lineage sizes *ℓ* (total offspring numbers of one stem cell) approaches a power-law:

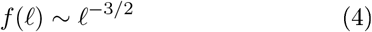

for late time points and large lineage sizes. We now argue that this is exactly what we expect if tissue growth is driven by a critical stem-cell population. Each stem cell (S-cell) population can be described by a time dependent trajectory *s* (*t*) which will start out with *s*(*t* = 0) = *s*_0_ cells and evolve until it eventually dies out at time *τ*: *s*(*t* = *τ*) = 0. We will call *τ* the trajectories *extinction time.* During each time interval *dt*, the *s*(*t*) S-cells will produce *g_D_s*(*t*)*dt* differentiated cells, and the total lineage size will be given by

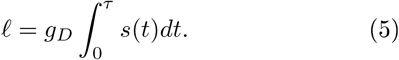

This equation is useful even without access to the precise trajectory *s*(*t*) because it allows us to compute approximations of ℓ by inserting any likely trajectory that leads from *s*(0) = *s*_0_ to *s*(*τ*) = 0. In section II F we will derive the most likely such trajectory from the path integral formulation of Brownian motion and use it to define ℓ as a function of *τ*. In particular, this will show that for large *τ*,

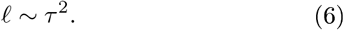

Since the extinction time *τ* is a random variable whose (cumulative) distribution function *p_D_*(*τ*) is given in equation (3), it follows as claimed that

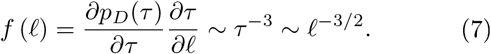

Without appealing to derivation in section II F, the scaling behavior ℓ ~ *τ*^2^ can be made plausible by comparing the evolution of the expected number of S-cells 〈*s*〉 from equation (2) to the extinction probability *p_D_* from (3). In the critical case, 〈*s*〉 = *s*_0_ holds, and the expected S-cell population size is thus constant. Yet every stem cell population certainly dies out eventually, and in particular we find (by cutting the exponential series in *p_D_* off after two terms) that the probability of a S-cell population surviving until a (large) time *t* scales like

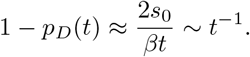

To keep the overall expected population size constant, it follows that S-cell populations that survive until time *t* must, on average, grow like *s*(*t*) ~ *t*[53]. The integration in equation (5) then yields, as anticipated

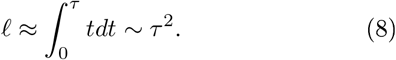

### C. Self organized criticality in experiments

The experimental data consists of lineage identifier counts for 15 organoids that were sequenced on days 16, 21, 25, 32 and 40 – three copies for each day[12]. Accounting for statistical and readout errors, the lineage identifier counts can be related to lineage sizes, i.e. to the numbers of cells in each lineage [43]. The organoids grow autonomously from day 11 onward, when they are not subjected to intrusive procedures anymore. Our model depends on four parameters which are determined from experimental data: *α* – the net growth rate of the S-cell population, *β*, measuring the stochasticity of the process, *s*_0_ – the average stem cell population per lineage at day 11 and *N* – the number of N-cells that are produced by each asymmetrically dividing A-cell. It is striking that the data seems roughly distributed according to an *x*^−3/2^ power-law (see Fig. 2). Therefore, we can assume that we are near criticality and *α* ≪ *β* holds. To determine the parameters precisely, we use the empirical characteristic function of the data

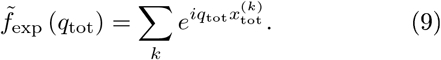

Here 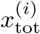 are the experimentally determined lineage sizes. The characteristic function of our model 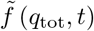 (see Supplementary Eq. (S 33)) is then least squares fitted to fexp 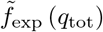. Moving the fitting procedure to Fourier space has several advantages: 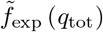 is less noisy than the empirical probability distribution and no smoothing procedures such as kernel density estimates need to be used. Finally approximations that are needed when transforming 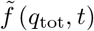 to real space are avoided. Our estimates for the model parameters are shown in Table I. It is interesting to visualize the estimates obtained by fitting the characteristic functions in terms of probability densities. In Fig. 3, a), we show a comparison between the probability densities of the SAN model *f*_tot_ (*x*_tot_, *t*) and the empirical probability densities at days 16 and 40. The empirical probability densities are obtained by taking the discrete derivatives of the cumulative probability distributions

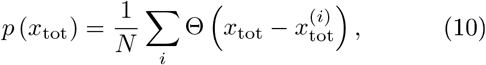

where *N* is the toW number of data points 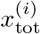 and Θ (*x*_tot_) is the Heaviside theta function with Θ (0) = 1. *f*_tot_ (*x*_tot_, *t*) is found using a Fast Fourier Transform of the characteristic function 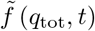.

**Table I.** Parameters of the SAN model estimated from experimental data with one standard deviation errors, *α* is the net growth rate of the stem cell population. The small value of *α* as compared to the stochasticity rate *β* indicates that the system is close to criticality, *s*_0_ and *N* are the average initial number of stem cells and the average number of N-cells produced by each stem cell, respectively.

|  |  |
| --- | --- |
| $\alpha$ [day<sup>-1</sup>] | $-0.018 \pm 0.004$ |
| $\beta$ [day<sup>-1</sup>] | $4 \pm 1$ |
| $s_0$ | $9 \pm 3$ |
| $N$ | $2 \pm 1$ |

The estimates of Table I show that organoid growth is indeed a self organized critical process with |*α*| ≪ *β*. It also shows that even at day 40 the process is far from its *t* → ∞ limit (*αt* ≈ 0.8) and the organoids maintain an avalanche-like population of S-cells at large *x*_tot_.

### D. Full Fokker-Planck description of lineage dynamics

Having shown how long-tailed power-law distributions can arise from critical proliferation and differentiation processes, we turn to a more detailed description of the biological processes and fix several drawbacks of the theory presented *s*_0_ far. We lift the assumption that the stem-cell population goes extinct and extend the Fokker-Planck equation (1) to include stem cell differentiation and asymmetrical divisions as separate processes. This allows us to develop a dynamical theory of tissue growth. Solving the Fokker-Planck equation, we obtain analytical expressions for the time dependent lineage size, which, at criticality, approaches the 3/2-power-law of Eq. (7) at long times. Modeling the system’s time dependence turns out to be crucial, since the in vitro data are taken at different stages of the organoids’ development, making it possible to observe how the lineage sizes approach the critical distribution.

In our model, the continuous growth and differentiation of a cell lineage consists of four processes: i) symmetrical S-cell division (S→S + S) at a rate *g_S_*, ii) S-cell differentiation into an A-cell (S→A) at a rate *g_A_*, iii) S-cell death (S→0) at a rate *g*_0_, iv) asymmetrical division of an A-cell into another A-cell and an N-cell (A→A + N) at a rate *g_N_*. and v) final differentiation of an A-cell into an N-cell (A→N) at rate *g_F_*. The rates and processes are summarized in Fig. 1. To describe these processes, we first have to write down a comprehensive master equation. However, the individual populations of S, A and N-cells are not accessible in lineage tracing experiments, since all descendants of a given stem cell inherit the same lineage identifier. To compare our theory to experiments, we therefore need to calculate the probability distribution of the total lineage size *x*_tot_ = *s* + *a* + *n*. This becomes simpler if we slightly modify our model by replacing the processes that generate N-cells (A→A + N and A→N) by simply assuming that each A-cell produces *N* = 1+ *g_N_/g_F_* N-cells over the course of its existence (where the +1 stems from the final A→N conversion) (Fig. 1). Since the N-cell output per A-cell varies by multiple orders of magnitude less than the total offspring of each S-cell[43] this simplification does not affect the main predictions of the model, but makes the model analytically tractable. Below we denote this process as S→(A)→ NN. The total lineage size becomes *x*_tot_ = *s* + *n*. With this assumption, the master equation for the probability distribution *f*(*s, a, n, t*) of S- and N-cell numbers *s, n* at time *t* reads

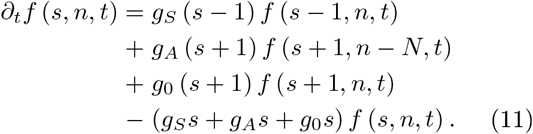

The right hand side terms of Eq. (11) correspond to the different processes that the cells undergo: Each combination of cell numbers *s, n* represents a state which the system might occupy with the probability *f* (*s, n, t*). The time derivative on the left hand side indicates that we are considering the rate of change of *f* (*s, n, t*). The terms *g_S_* (*s* – 1) *f* (*s* – 1, *n, t*) and −*g_S_sf*(*s, n, t*) describe symmetric S-cell divisions at a rate *g_S_*. Stem cell death is described by terms proportional to *g*_0_. The second right hand side term accounts for an S→(A)→ NN transition while the system is in a state with *s* + 1 S-cells. The term −*g_A_sf* (*s, n, t*) represents an S→(A)→ NN process with the system being in a state with *s* S-cells. We translate the discrete process given by (11) into a continuous version given by the Fokker-Planck equation [54]:

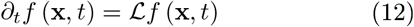

with the differential operator

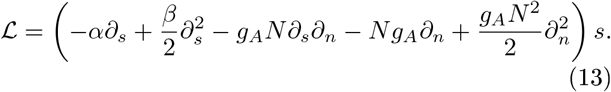

Here, **x** = (*s,n*) is a vector with the cell numbers as *α* = *g_S_* – *g_A_* – *g*_0_ and *β* = *g_S_* + *g_A_* + *g*_0_ *s* and *n* again become continuous variables, and as for the pure S-cell process we may interpret the fractional part as reflecting the cells internal states. In fact, the first two terms of Eq. (13) describe the same S-cell dynamics as the pure S-cell process from equation Eq. (1). The remaining terms describe cell differentiation and asymmetrical division.

### E. Tissue growth: power-laws and avalanches

Next, we want to examine the implications of the Fokker-Planck Eq. (12) for the lineage sizes within a tissue sample. To this end we solve Eq. (12) with the initial condition *f* (*x, t* = 0) = *δ* (*s* – *s*_0_) *δ* (*n*). Using the Fourier transform of Eqs. (12), (13) with respect to **x** and the method of characteristics to solve the resulting first order partial differential equation (see Supplementary Sec. A), we find the characteristic function of *f* (**x**, *t*), 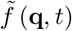, which is it’s Fourier transform: 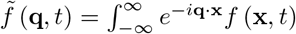.

Since experiments measure the total lineage size *x*_tot_ = *s* + *n* and not the individual cell numbers *s* and *n*, we are interested in the probability distribution for *x*_tot_, which we call *f*_tot_ (*x*_tot_,*t*). This distribution is given by an integral of *f* (**x**, *t*) over all states with equal *x*_tot_:

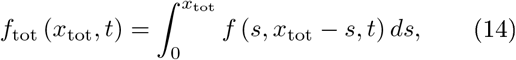

where *f* (**x**, *t*) is the solution to Eq. (12).

We find that the behavior of *f*_tot_ depends on the sign and value of the S-cell growth rate *α*. For large times *t* → ∞, *f*_tot_ (*x*_tot_, *t*) approaches a limiting distribution 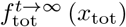, which is a truncated 3/2-power law:

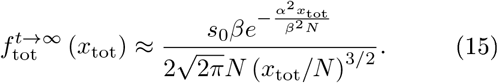

At criticality (*α* = 0), the exponential truncation vanishes, and the distribution becomes a true 3/2-power-law. For small |*α*|, the truncation will only become important at very large *x*_tot_/*N* > *β*^2^/*α*^2^. In other words, the smaller the growth rate, the more pronounced the power-law.

The way in which *f*_tot_ (*x*_tot_, *t*) approaches the limiting distribution of Eq. (15) is very different for *α* > 0 and *α* < 0. For positive *α*, for any finite *t* and large enough lineage size 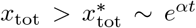, there is a regime where *f*_tot_ (*x*_tot_, *t*) is not approximated by Eq. (15). This is the *avalanche regime* illustrated in Fig. 1. Here lineages have a high percentage of S-cells which are proliferating and driving the system towards larger lineage sizes, while for 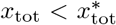 most lineages are fully differentiated and do not grow any more. While the size of lineages in the avalanche regime can become very large, the probability with which such large lineage sizes occur becomes smaller and smaller. This behavior is illustrated in Fig. (3) c). In the avalanche regime, *f*_tot_ (*x*_tot_, *t*) can be approximated by

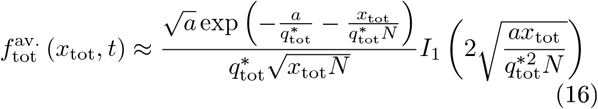

where *I*_1_ (*z*) is the modified Bessel function of the first kind and 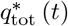 and *a* (*t*) are non trivial time dependent coefficients which are defined in Supplementary Eqs. (S 37) and (S 42). This result is derived in Supplementary Sec. A.

For a negative *α*, we also find an avalanche regime hosting an active S-cell population (even though the avalanche will stop eventually). This regime is located at large *x*_tot_, and is marked by a region where *f*_tot_ (*x*_tot_, *t*) is larger than the limiting distribution 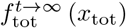, followed by a rapid truncation at even larger *x*_tot_ (Fig. 3 b)). In contrast to the behavior at *α* > 0 *f*_tot_ (*x*_tot_, *t*) convergences to 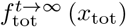 uniformly, i.e. avalanches becomes less and less pronounced as *t* → ∞. This is related to the fact that all lineages eventually die out if *α* < 0 [52]. Our estimates show, that *α* ≲ 0 holds in experiments (see Sec. II C). The analytical approximation (16) holds for the avalanche part of the lineage size distribution for *α* < 0, if *t* is sufficiently small, i.e. *αt* ≲ 1. In particular, it holds reasonably well for the experimental data, as we demonstrate in Fig. 3 b). This has to do with the analytical structure of the characteristic function 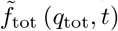 which is discussed Supplementary Sec. A.

### F. Extinction Trajectories

We have shown in section II E that the appearance of a 3/2 power law in the distribution of lineage sizes in the SAN model is intrinsically linked to criticality, i.e. to a vanishing S-cell net grow rate *α*. Here we develop a path integral approach to the SAN-model based on the Onsager-Machlup formalism [55, 56]. This allows us to study the trajectories of individual lineages *s* (*t*) and the emergence of the 3/2 power-law from the point of view of possible fates of a single S-cell lineage. In particular, we find that critical trajectories can be divided into three qualitatively distinct classes depending on how long-lived the population is.

As in Eq. (1) and the following discussion, we restrict ourselves to S-cells only and extend the results to capture the full SAN model later on. For the critical case *α* = 0, equation (3) stated the probability (*t*) = *e*^−2*s*_0_/*βt*^ that a lineage initially consisting of *s*_0_ S-cells has died out by time *t*. This probability tends towards one as *t* goes to infinity, and every lineage’s S-cell population thus dies out eventually. Consequently, every sample path **x**(*t*) = (*s*(*t*), *n*(*t*)) of the diffusion process described by (13) has a well defined time *τ* at which *s*(*t*) first reaches zero, we call this the path’s *S-cell extinction time τ*. Among the many different paths that a lineage’s S-cell population can take given its initial size *s*(0) = *s*_0_ and extinction time *τ*, we call the most likely path *s_τ_* (*t*) the *S-cell extinction trajectory.* To find *s_τ_* (*t*), it is convenient to transform the Fokker-Planck equation (1) into a stochastic differential equation for the variable *s*:

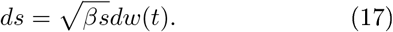

Here *w*(*t*) is the standard Brownian motion. Introducing the new variable 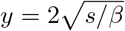, Eq. (17) is written as

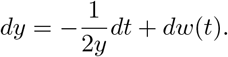

In contrast to Eq. (17), the coefficient in front of the Brownian motion term is constant in the above equation, and hence a Lagrangian 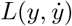 can be constructed [55, 56]. The minimization of the corresponding action 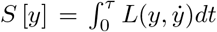 under the constraints of the initial and final boundary conditions (*s*_0_ initial cells and extinction at time *τ*) leads to an optimal path *y* (*t*). Transforming back, we find the extinction trajectory *s_τ_* (*t*):

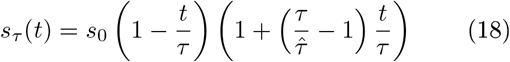

with a characteristic time scale 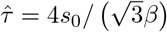. A more complete description of the path integral approach can be found in Supplementary Sec. B.

The shape of the extinction trajectory *s_τ_* (*t*) depends on how far the extinction time *τ* lies in the future compared to the characteristic time 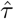. For 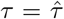, the lineage size will decrease linearly: 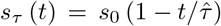. In general we can find that the number of S-cells will likely

1. decrease faster than linearly if 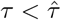,
2. decrease slower than linearly if 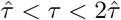 (exactly linear if 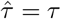),
3. and initially grow if 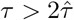.

Figure (4) shows the possible shapes of S-cell extinction trajectories, and shows that the analytically derived *s_τ_* (*t*) agrees with numerically estimated extinction trajectories of both the continuous S-cell process from equation (1) as well as for its discrete counterpart. Crucially, the result of Eq. (18) shows that in the 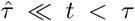 limit, the trajectory 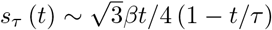 does not depend on the initial number of cells *s*_0_ (see Fig. 4). Showing that at this point the lineage lost all memory of its initial size.

We now return to the full SAN process from equation (13). If a lineage’s S-cell population follows the extinction trajectory *s_τ_* (*t*), we expect it to produce A-cells with rate *g_A_s_τ_*(*t*) at time *t*, and expect each produced A-cell to produce *N* = 1 + *g_N_/g_F_* N-cells over the course of its existence (see the discussion leading up to equation (11)). Once a lineage’s S-cell population goes extinct at time *τ*, the total lineage size will thus approximately be

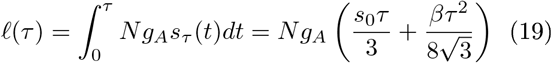

This delivers the functional relationship between a lineage’s S-cell extinction time and its final size. This relationship again shows that the effect of the initial size *s*_0_ on a lineage’s eventual total size (*τ*) diminishes as the lifetime *τ* of its S-cell population increases. While for short-lived S-cell populations the final total lineage size depends linearly on *s*_0_, for long-lived S-cell populations *ℓ*(*τ*) doesn’t scaled with *s*_0_, but instead scales quadratically with *τ*, as was already anticipated in equation (6).

The 3/2 power-law governing the size distribution of (large) lineages under criticality then emerges from the asymptotic scaling law *τ*(*τ*) ~ *τ*^2^ as outlined in Sec. II B. For a lineage to eventually reach size *x*_tot_ its extinction time must be *τ* ≈ ℓ^1^ (*x*_tot_) where *τ* has probability density *dp_D_* (*τ*)/*dτ*. For the probability density of *x*_tot_ this yields

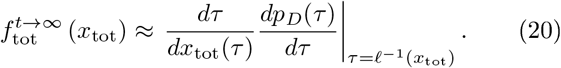

Since these quantities scale like 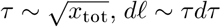 and *dp_D_* (*τ*) ~ *τ*^−2^*dτ*, in the limit of large lineage sizes (and thus large extinction times) we get

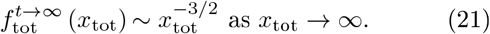

From the path integral perspective, we have approximated the total population size *x*_tot_ averaged over all paths satisfying *s* (0) = *s*_0_ and *s* (*τ*) = 0 by replacing the average with the most likely path. The remaining random variable is *τ* with it’s probability density *pD* (*τ*). In Eq. (20) we then changed the variables from *τ* to *x*_tot_. In the spirit of this approximation, we can also obtain the numbers of A- and N-cells through the rate equations

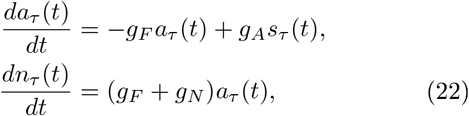

which can be derived from the master equation (11), where *s_τ_* (*t*) is given by Eq. (18). For a fixed time *t*, the distributions *f*_s_(*t*|*τ*) of *s_τ_*(*t*), *f_n_*(*t*|*τ*) of *a_τ_*(*t*) and *f_α_*(*t*|*τ*) of *n_τ_*(*t*) can then be found as before by considering these to be functions of *τ* which has a known cumulative distribution function pD. Finally, the total lineage size *x*_*t*0_ *t*(*t*|*τ*) of a lineage whose S-cell extinction time is *τ* is then *x*_tot_(*t*|*τ*) = *s_τ_* (*t*) + *a_τ_* (*t*) + *n_τ_* (*t*), and its (approximate) distribution can be found as in equation (20) by a change of variables from *τ* to *x*_tot_. It is helpful to note that *s_τ_* (*t*) and hence also *a_τ_* (*t*), *n_τ_* (*t*) and *x*_tot_(*t*) increase monotonically with *τ* for any fixed time *t*, and *x*_tot_(*t*|*τ*) is therefore a *one-to-one* mapping between S-cell extinction time *τ* and total lineage size at fixed time t. Thus extinction trajectories offer an alternative approach to calculating the probability densities at criticality.

## III. DISCUSSION

We introduced the SAN model to describe the growth of human cerebral organoids and demonstrated the critical nature of the process. Considering the small number of parameters (see Table I) of the model, our theory fits well to the lineage tracing data of Ref. [12].

We pointed out that the criticality of the process implies the presence of a mechanism tuning the rates of division and differentiation to criticality. Different mechanisms that balance stem cell division and differentiation are known, and can be classified according to their scope as either (a) balancing on the level of single cells, or (b) balancing on the population level [2]. Balancing at the level of single cells implies a form of asymmetric division in which only one daughter cell of a stem cell inherits the stemness property, while the other proceeds to differentiate. This mechanism, while perfectly balancing differentiation and division, is deterministic and therefore cannot explain the observed variety in lineage sizes. We conclude that divisions and differentiations are balanced at the population level instead, either through chemical communication between cells, or by competition for a scarce resource such as space or nutrients. While characterizing the regulatory mechanisms involved with any certainty would require more data, comparison with other biological systems suggests the niche mechanism as a possibility. In the mechanism, found, among others in intestinal crypts and in the adult human brain, stem cells are located in niches in which the available space restricts the number of possible divisions: with each symmetric division of a stem cell, the containing niche becomes more crowded which increases the chance of a stem cells being pushed out of the niche, thus triggering its differentiation [7, 57, 58].

Despite being surprisingly robust, the SAN model is only a minimal model of biological reality and as such cannot account for the full spread of the experimental data.For example, the model predictions and the empirical distributions differ for small *x*_tot_ (see Fig. 3). This is mainly due to two reasons: First, many small lineages have died out in the early stages of the organoid development. Dead lineages are obviously not described by SAN dynamics. Second, deriving the analytical results we assumed that all lineages consist of *s*_0_ stem cells at day 11. While we determine *s*_0_ from the data, it is clear that the value is only an average over the different lineages. While for the very large, lucky lineages the initial number of stem cells is not important, it matters for lineages that differentiated quickly and did not have the opportunity to grow a large number of S-cells.This is a crucial aspect of critical, stochasticity driven growth (see Sec. II F). Furthermore the time dependence of division and differentiation rates is not captured by the SAN model. Both the variability of *s*_0_ as well as the time dependencies of rates contribute to the fact that the actual spread of the empirical data goes beyond what is predicted by the Fokker-Planck model. Another source of inaccuracy may be the implicit assumption that the division and differentiation processes are Markovian. In principle, cell division is not memoryless, but instead the result of a sequence of cell state changes (G1→S→G2→M). Moreover our simplifying assumption that each A-cell produces *N* N-cells at a rate *g_A_* is valid only if the time intervals between observations are large enough.

Note that the model uses one parameter set (Table I) to describe not only the overall shape of the probability densities, but also their time dependence. In particular, the time dependence of the truncation for large lineage sizes is well captured (Fig. 3 a)). Support for our model also comes from an extensive computational experiment [43] aimed at estimating the rates of division and differentiation processes in human cerebral organoids using the same data set. With our choice of parameters, Ref. [43] reports *β* = 3.37, *α* = —0.01 with large uncertainties that have the same order of magnitude as the results. While the biological modeling assumptions of our work and Ref. [43] are similar, the fitting procedure and data handling are completely independent. The fact that the rate estimates are in good agreement is a strong argument supporting the validity of the Fokker-Planck approach of Eq. (12) and the conclusions that are drawn in this work.

In summary, this paper presents strong evidence that human cerebral organoid growth is a self organized critical process, and that cells tune themselves to the point of maximal stochasticity, avoiding the archetypal scenario of exponential growth and arrest. As in the case of simple sand piles, maintaining the critical state requires a mechanism through which individual cells become aware of the current balance between division and differentiation events in the organoid. The search for such a mechanism could inspire future experimental research. Recent advances in lineage tracing allow to study the spatial distribution of lineages in cerebral organoids using lightsheet microscopy and spatial transcriptome sequencing [59]. Inferring the spatial dynamics of the growth process could help to identify the the mechanism behind organoid SOC. Quantitative lineage tracing experiments with other organoid types such as intestinal [60, 61] or cardiac [62, 63] organoids could reveal whether critical growth is specific to cerebral tissue, or whether it is a more general organizing principle.

## ACKNOWLEDGMENTS

We thank C. Esk, S. Haendeler, B. Jeevanesan, J. F. Karcher, and D. Lindenhoferfor inspiring discussions. This project has received funding from the European Union’s Framework Programme for Research and Innovation Horizon 2020 (2014-2020) under the Marie Curie Skłodowska Grant Agreement Nr. 847548 (AvH, EK), and from a Special Research Programme (SFB) of the Austrian Science Fund (FWF), project number F78 P11 (AvH, FP).

## SUPPLEMENTARY MATERIAL

### A. Solving the Fokker-Planck equation

#### 1. The characteristic function

The Fokker-Planck equation (12) can be solved in the Fourier domain. After a Fourier transform, Eq. (12) becomes

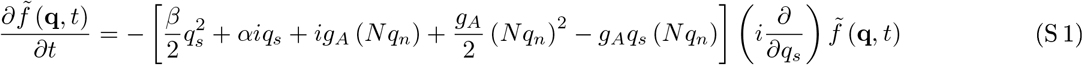

with

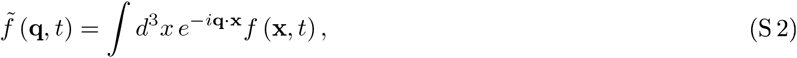

where **q** = (*s, n*). In the following we will drop the factors of *N* for notational simplicity and restore them later on with the substitution

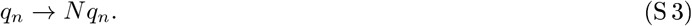

Let us choose the initial condition

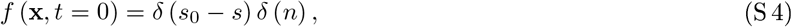

which corresponds to *s*_0_ S-cells at *t* = 0. In Fourier space this initial condition translates to

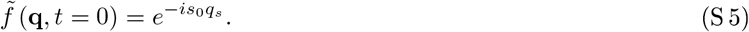

Eq. (S 1) contains only first order derivatives and can be solved with the method of characteristics. The characteristic equations are

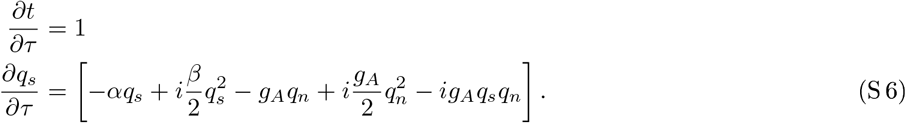

The first equation is trivial. It is solved by *t* = *τ*. The second equation is solved by

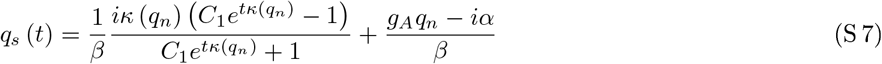

with

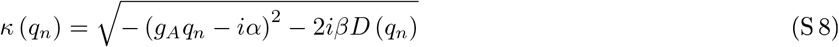

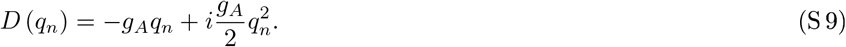

The solution for the Fourier transformed Fokker-Planck equation (S 1) is found by solving for the integration constant *C*_1_:

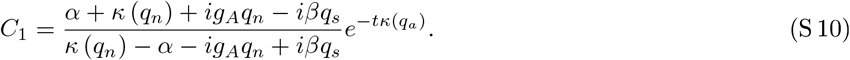

The next step is to look for a function of *C*_1_, *G* (*C*_1_ (*q_s_, q_n_, t*)), which satisfies

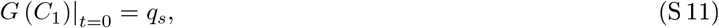

so that a function 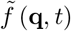 that satisfies the initial condition of Eq. (S 5) can be written as

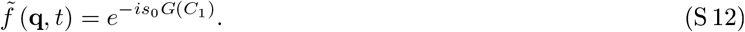

Such a function is given by

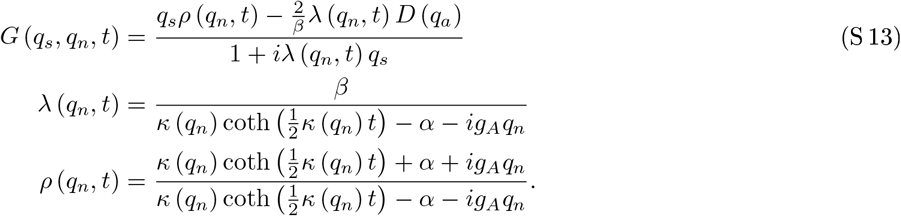

The function 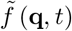 of Eq. (S 12) is the characteristic function of *f* (*x, t*). For later purposes we want to separate *κ* (*q_n_*) into real and imaginary parts. We find

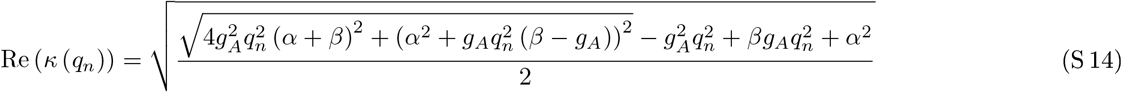

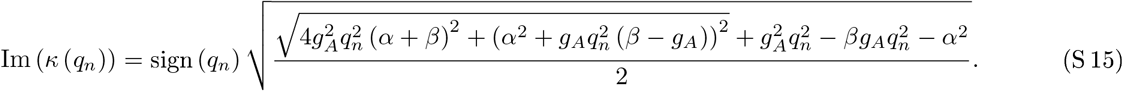

#### 2. Large t limit. Distribution of N cells

Let us gain some intuition into the dynamics of the growth process. If the S-cells are dividing at a near critical near zero rate *α*, while they are differentiating at a much larger rate *g_A_* we can expect that, as *t* grows, most lineages will consist of differentiated cells. Lineages that escaped differentiation will have to be very lucky and sufficiently large. As a first step, we want to consider the distribution differentiated cells.

The inverse Fourier transform of Eq. (S 12) with respect to *q_s_* reads

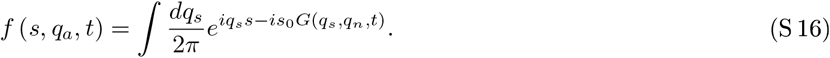

This expression can be rewritten as

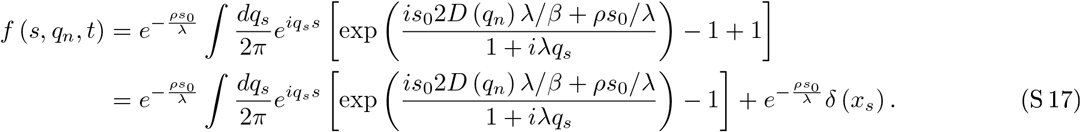

We have thus separated the function *f* (*x_s_, q_n_, t*) into a part which depends on *x_s_* and describes lineages that are still evolving and a second part with *x_s_* = 0. This latter part contains information on the distribution of N-cells in fully differentiated lineages. Its inverse Fourier transform with respect to *q_n_* is given by

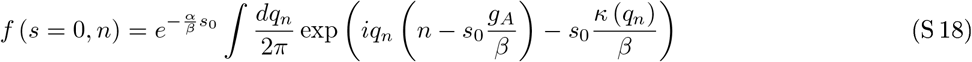

Notice that *f* (*s* = 0, *n*) does not depend on time. We will later see that *f* (*s* = 0, *n*) is the *t* → ∞ limit of the distribution of total lineage sizes (see Eq. (14) in the main text). Note that the integrand of the first right hand side term in the second line of Eq. (S 17) approaches zero as *q_s_* → ∞. This is a consequence of separating out the delta function and is necessary to regularize the integral. We will attend to this issue below. To do the integral in Eq. (S 18), it is useful to take a look at the analytic structure of the integrand. Nonanalyticities arise from the square root structure of *κ* (*q_n_*). Two branch cuts start at the two imaginary roots of *κ* (*q_n_*) and run to +*i*∞ and – *i*∞ along the imaginary *q_n_* axis (see Fig. S 1). The positive imaginary root is

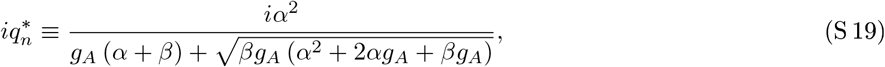

while for the negative imaginary root we find

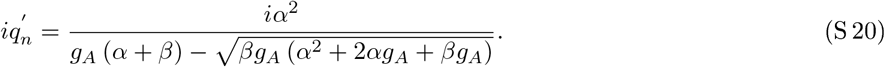

**Supplementary Figure S 1.**
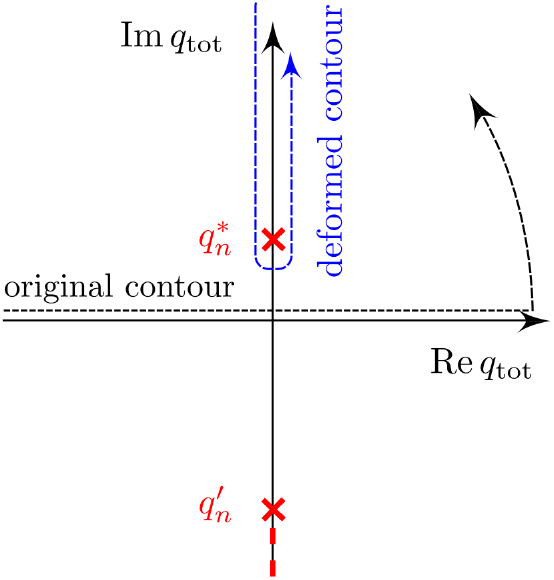
The branch-cut and contour integration in Eq. (S 23).

For small |*α*| ≪ *β*, we have

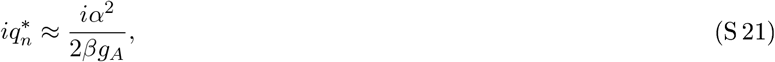

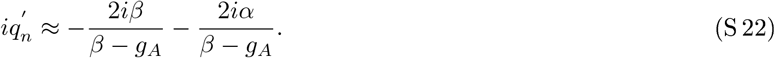

We thus make a crucial observation: The branch-cut in the upper complex half-plane of *q_n_* descends to the origin for *α* = 0, while the branch cut in the lower half plane always starts below – 2*iβ*/ (*β* – *g_A_*). For *n* > 0, the integral over the real line in the inverse Fourier transform in Eq. (S 18) can be deformed to an integral around the branch cut running from +*i*∞ to 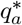 to the left hand side of the branch cut (at a small distance *ε*, say), and then running back to infinity on the right hand side (see Fig. S 1). The half-circle around 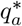 is of order *ε* and can be neglected. We can write

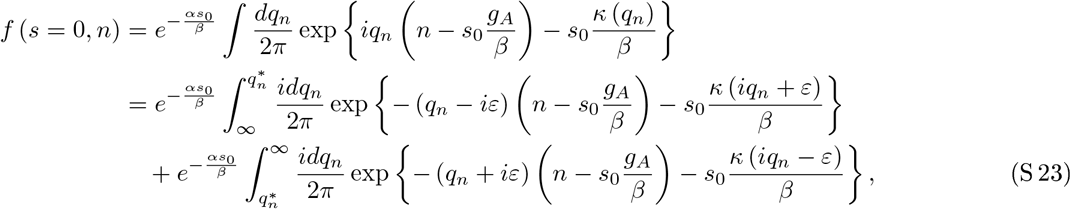

On the new contour, we need

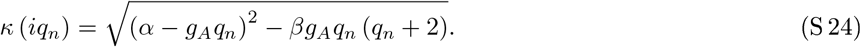

It is clear from Eq. (S 23) that the most important contribution to the integrals will come from the vicinity of 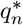, because the integrand decays exponentially for larger *q_n_* the faster, the larger *n*. Expanding *κ* (*iq_n_*) around 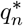, we obtain

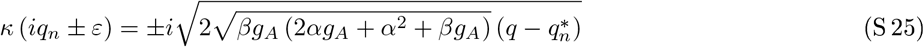

(approaching the branch cut from different sides changes the sign of the square root). Thus, for large *n*, we approximate Eq. (S 23) as

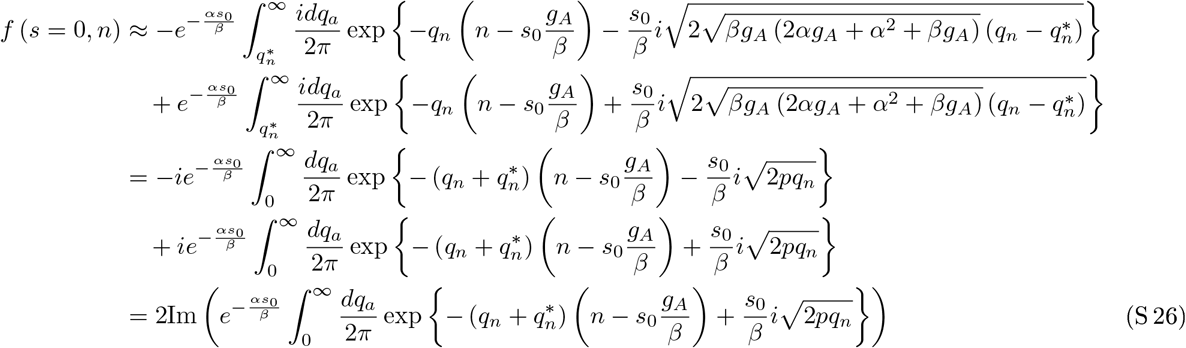

with

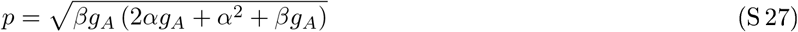

For the remaining integral we obtain

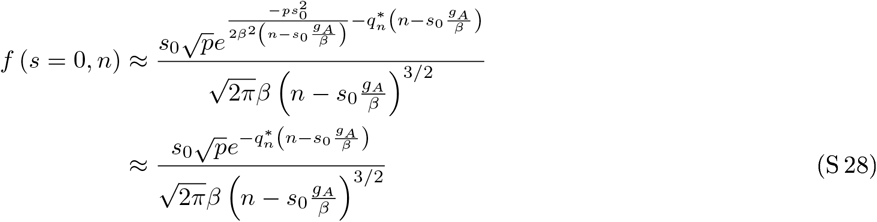

This is a truncated, one-sided Lévy distribution [64–66]. The appearance of the Lévy distribution could have been anticipated, since the integrand of Eq. (S 18) for *α* = 0 can be mapped to the characteristic function of the Lévy distribution after the argument of the square root in *κ* (*q_n_*) lias been expanded for small *q_n_*. For small *α* and *g*_0_ we find *g_A_* = (–*α* + *β* – *g*_0_) /2 ≈ *β*/2, as well as *p* ≈ *β*^2^/2, and for *n* ≫ 1 the truncated power law of Eq. (S 28) becomes Eq. (15) of the main text with *N* = 1. In the main text, we stated that the *t* → ∞ limit of *f*_tot_ (*x*_tot_, *t*) – the distribution of total lineage sizes *x*_tot_ – is governed by the same expression as *f* (*s* = 0, *n*) (see Eq. (15) of the main text). We will justify this statement below. On an intuitive level this can be understood as a consequence of the fact that, as *t* increases, most cells will already have differentiated and *f*_tot_ will be dominated by the N-cell distribution.

Since 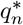 descends to the origin as *α* → 0, the truncation of the 3/2-power-law of (S 28) vanishes and (S 28) becomes a true Lévy stable power-law distribution. Returning to the observation that the integrand of (S 18) is nonanalytic in the lower complex half plane, we conclude that *f* (*s* = 0, *n*) is finite for *n* < 0, meaning that there is a finite probability to find a negative number of N-cells. This is a consequence of approximating the discrete master equation by a continuous Fokker-Planck equation which is accurate for large *s* and *n*. However, one can easily convince oneself that *f* (*s* = 0, *n*) decays very fast as *n* decreases below zero: Following the reasoning of the above branch-cut integration, we see that *f* (*s* = 0, *n*) will be suppressed by an exponential factor 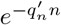. On the other hand, we see from Eq. (S 22) that for small *α*

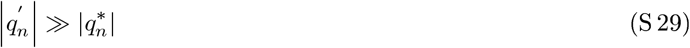

holds. Thus we conclude that the probability for a negative *n* is very small, and can be neglected as an artifact of the Fokker-planck approximation. Notice that the probability for negative *s* is zero, because Eq. (S 12) is analytic in *q_s_*, except for a single pole which is always in the upper complex half plane (since Re (λ (*q_n_*) > 0) as Eqs. (S 14), (S 13) indicate). Indeed, while the diffusion coefficient associated with the second derivative in *s* in Eq. (13) vanishes at *s* = 0, the diffusion coefficient associated with *n*-diffusion does not vanish at the origin and allows for a small leakage of probability towards negative *n*.

So far we have been investigating the analytic structure of the *s* = 0 contribution to 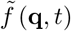. However similar arguments carry over to the general case. Multiplying the numerator and denominator of the function *G* (*q_s_, q_n_, t*) in Eq. (S 13) by λ (*q_n_, t*)^−1^ and keeping *q_s_* reffi for the moment, we see, that the characteristic function 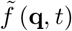 (Eq. (S 12)) is analytic in *q_n_* except for the branch-cuts of *κ* (*q_n_*) and the pole at λ (*q_n_, t*) = *i/q_s_*. This pole occurs when λ (*q_n_, t*) is purely imaginary, *f* (*s, n, t*) will decay for *n* < 0 the faster, the lower the nonanalyticities with Im (*q_n_*) < 0 lie in the complex plane. Eq. (S 29) shows that the contribution of the lower half branch-cut is indeed negligible since it starts much further away from the origin than the upper branch-cut. A numerical inspection of λ (*q_n_, t*) shows that regions where λ (*q_n_, t*) is purely imaginary in the lower complex half plane of *q_n_* are indeed sufficiently far from the origin for all reasonable parameter choices. We conclude that the finite values of *f* (*s, n, t*) for negative *n* are simply an artifact of the Fokker-Planck approximation to the original master equation (11) and are small.

#### 3. *The lineage size distribution f*_tot_ (*x*_tot_, *t*)

As pointed out in the main text, we are ultimately interested in the distribution of the lineage size

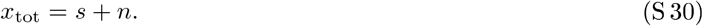

This distribution is obtained by summing *f* (*s, n, t*) over all states which satisfy the condition (S 30):

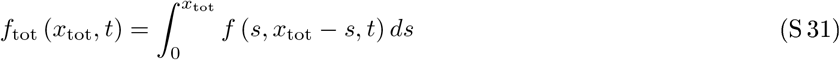

(see also Eq. (14) in the main text). Having argued in Sec. Supplementary Sec. A 2, that the distribution *f* (*s, n, t*) corresponding to the characteristic function (S 12) is confined to positive *s* and – to a good approximation – to positive *n*, we can extend the integration in Eq. (S 31) to the complete real axis:

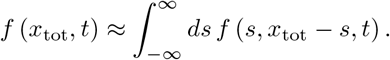

A Fourier transform then gives

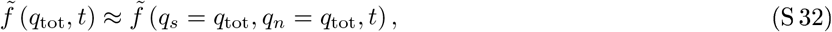

where the characteristic function of Eq. (S 12) is on the right hand side. This yields

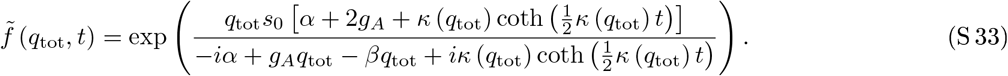

##### Analytic expressions for the power-law and avalanche regimes

The characteristic function given in Eq. (S 33) has a peculiar behavior at *q*_tot_ = 0 which determines the asymptotics for large *x*_tot_. To see this, let us first investigate the *t* → ∞ and the *q*_tot_ → ∞ limits of 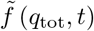. Since Re*κ* (*q*_tot_) > 0 holds for all real *q*_tot_ (see Eq. (S 14)) and Re*κ* (*q*_tot_) ~ |*q*_tot_| for large arguments, the approximation

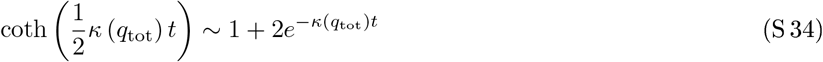

holds in both, the *t* → ∞ and the *q*_tot_ → ∞ limits. Using (S 34), Eq. (S 33) can be approximated by

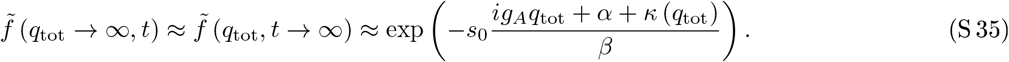

For *α* > 0, this expression approaches exp(−2*αs*_0_/*β*) as *q*_tot_ → 0, whereas for *α* ≤ 0, it approaches unity. For the full characteristic function, however, 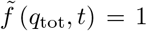 is true in all cases. This is depicted in Fig. S 2. We conclude that Eq. (S 35) is not a good approximation for small |*q*_tot_| if *α* > 0 bolds. To find an approximation for small |*q*_tot_ | and positive growth rates, we expand the numerator and denominator inside the exponential function in Eq. (S 33) around *q*_tot_ ≈ 0 and find

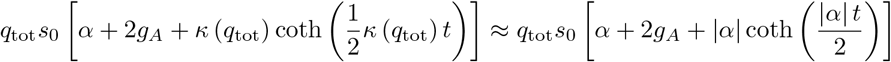

and

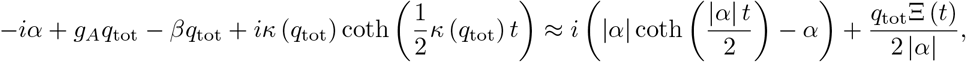

where

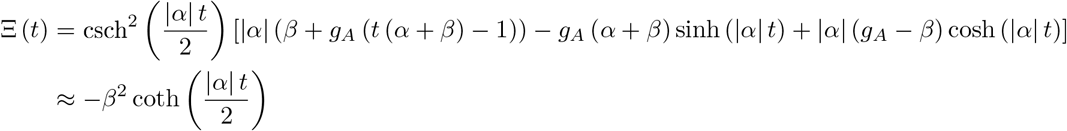

**Supplementary Figure S 2.**
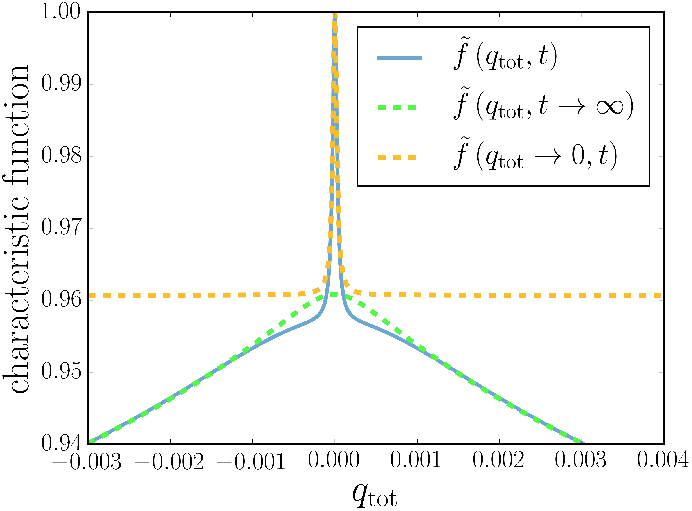
The full characteristic function 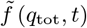 of Eq. (S 33) and the approximations for *t* → ∞ or large *q*_tot_, 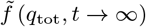, (Eq. (S 35)), as well as for small *q*_tot_, 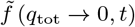, (Eq.S36). We used a positive growth rate *α* = 0.2/day and *β* = 10/day. For *α* > 0 and large but finite *t*, 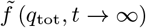 (green dashed curve) is a good approximation to the characteristic function 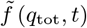 (blue curve), except for a small region near *q*_tot_ = 0 which is well approximated by 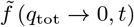 (orange curve). This region governs the behavior at large lineage sizes *x*_tot_.

In the last line we assumed that |*α*| ≪ *β*. Therefore, the appropriate approximation for *α* > 0 and small |*q*_tot_| reads

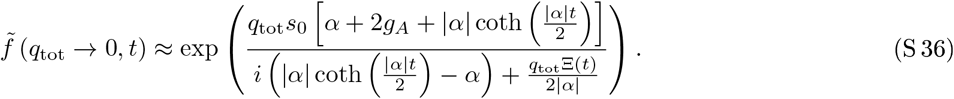

It remains to determine the distribution function *f* (*x*_tot_, *t*). At small and intermediate *x*_tot_, we expect *f* (*x*_tot_, *t*) to be governed by the behavior of the characteristic function at larger *q*_tot_, whereas the behavior of *f* (*x*_tot_, *t*) for large *x*_tot_ will be determined by the region around *q*_tot_ ≈ 0. This has to do with the analytic structure of Eq. (S 33). The question is, which nonanalyticity dominates the Fourier transform as the integration contour is deformed according to Fig. S 1. Besides the branch-cuts of *κ* (*q_n_*), 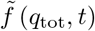 exhibits a pole on the imaginary axis (see Fig. S 3). Let this pole be located at 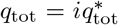. If it is closer to the origin than the upper starting point of the branch-cut, i.e. 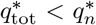, where 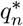 is defined in Eq. (S 19), the pole becomes dominant at large *x*_tot_. This is because both nonanalyticities, the branch cut starting at 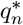 and the pole at 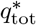, are located on the positive imaginary axis. Hence, when the integration along the deformed contour is performed, the contributions of the pole and the branch-cut will roughly behave as 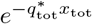 or 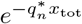, respectively. We can use Eq. (S 36) to track the behavior of 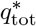 for different parameter values:

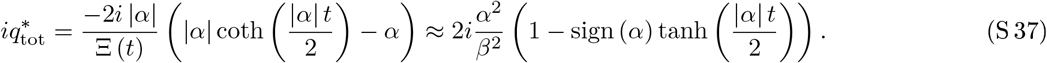

Eq. (S 37) shows that the analytic structure of the characteristic function is very different for positive and negative *α*. For positive *α*, it is 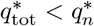, and therefore *f* (*x*_tot_, *t*) will be dominated by the single pole that is captured in Eq. (S 37). For *α* < 0, the opposite scenario is true: we find 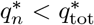.

We now calculate the inverse Fourier transforms Eqs. (S 35) and (S 36) which will give us approximate expressions for *f* (*x*_tot_, *t*) at intermediate and large *x*_tot_. The inverse Fourier transform of Eq. (S 35) reads

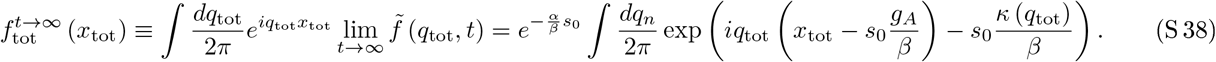

We encountered this integral when we calculated the distribution of N-cells in Eq. (S 18). Similarly to Eq. (S 28) we find

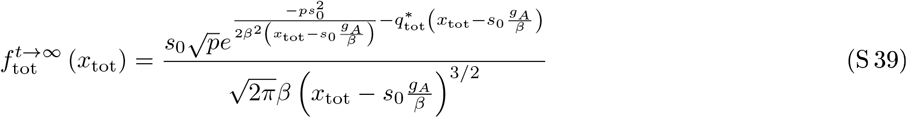

with

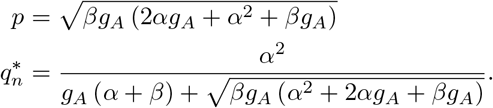

**Supplementary Figure S 3.**
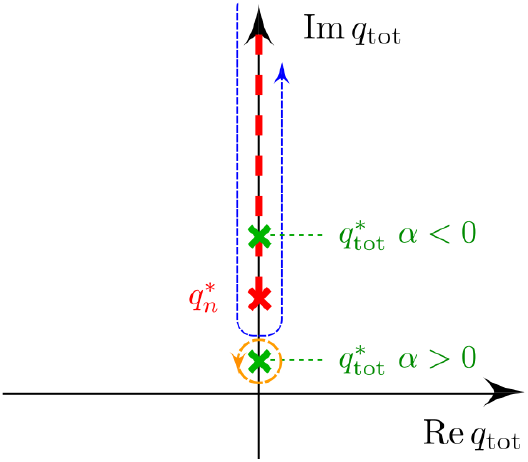
Branch-cut integration (see Fig. S 1 and Eq. (S 23)) and the pole at 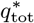 (Eq. (S 37)) characterizing the avalanche part of the distribution. Both, the pole and the branch-cut must be captured by the deformed contour (blue dashed line). For *α* > 0, the pole at 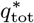 is always nearer to the origin than the starting point of the branch-cut 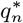 and thus determines the large *x*_tot_ behavior. For *α* < 0, 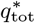 is within the integration contour, moving towards larger imaginary values as *t* is increased. For sufficiently small *t*, the avanlanche part of the distribution is still well approximated by the integration around the 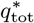 pole (see Fig. 3 b)).

In order to account for the *N* N-cells that each S-cell is producing we have to make the substitution *q_n_* → *Nq_n_* in Eq. (S 32). This substitution carries over to Eq. (S 35) where we have to replace *q*_tot_ by *Nq*_tot_. The integration of Eq. (S 38) is evaluated according to

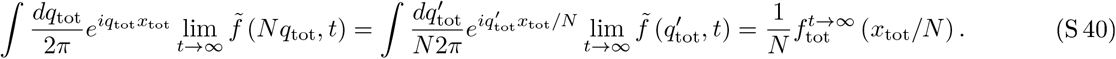

For |*α*| ≪ *β*, we obtain the result of Eq. (15), Sec. II E of the main text:

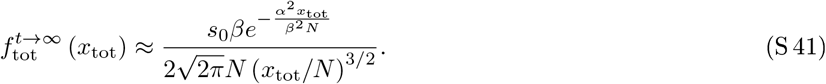

According to the above discussion, Eq. (S 41) is valid for intermediate *x*_tot_ if *α* > 0. For *α* < 0, the contribution of the pole becomes stronger and stronger sub-leading to the branch-cut contribution as *t* is increased (see Eq. (S 37)). However, if *t* is sufficiently smffil, the pole is in the vicinity of the branch-cut starting point 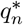 and has a strong influence on the distribution function. For large enough *t* however, Eq. (S 41) is a good approximation for *f*_tot_ (*x*_tot_) everywhere.

To determine the behavior at large *x*_tot_ for *α* > 0 we make use of Eq. (S 36) which gives a good approximation for the characteristic function at small *q*_tot_. It is useful to rewrite Eq. (S 41) as

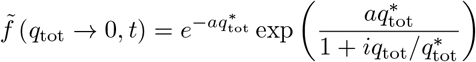

with

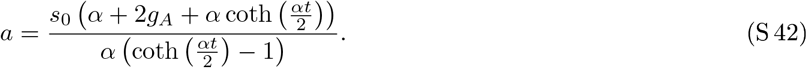

The inverse Fourier transform

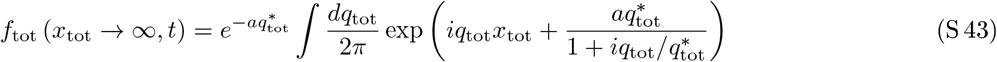

is strictly speaking divergent (as is the one in Eq. (S 17)). However it can be regularized by adding a small exponential factor *e*^−*q*_tot_|*ε*^ to the integrand and letting *ε* → 0 after the integral is done. The large *x*_tot_ behavior that we are interested in is dominated by the pole at 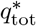 for *α* > 0. We will see later, that for *α* < 0, the pole still dominates the distribution for reasonably large *x*_tot_ (avalanche regime) if *t* is small, although it does not govern the asymptotics for *x*_tot_ → ∞. The pole’s contribution can be found by integrating along a circle around 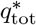 (see Fig. S 3). Writing

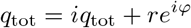

the integral around the pole at *q*_tot_ becomes

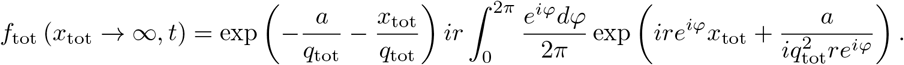

Since the radius *r* is arbitrary, we are free to chose

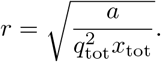

After some algebra, we arrive at

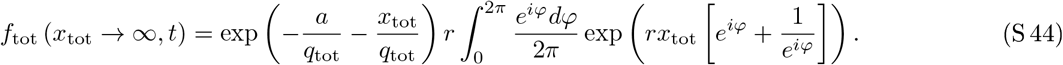

Using the integral representation of Bessel functions

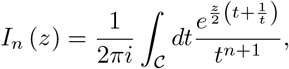

where 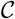 is a contour enclosing the origin, with the substitution *t* = *e^τψ^* Eq. (S 44) becomes

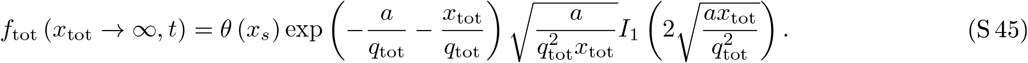

For *α* > 0, this is the large *x*_tot_ approximation for the distribution function *f* (*x*_tot_, *t*) given in Eq. (16), where we have restored the variable *N*. As is demonstrated in Fig. (3), Eq. (S 45) provides a good approximation for the large *x*_tot_ behavior of *f*_tot_ (*x*_tot_, *t*) for a positive S-cell growth rate *α*. However, even for *α* < 0. at reasonably large *x*_tot_ (namely in the avalanche regime), the behavior of the lineage size probability density is well described by Eq. (S 45), if *αt* ≲ 1 holds. This is due to the fact that the pole at 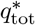, while located above 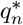 on the imaginary *q*_tot_ line (see Fig. S 3), is still sufficiently near 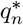 to dominate the avalanche part of the distribution function (see Eq. (S 37)). However, the asymptotics of *f*_tot_ (*x*_tot_, *t*) for *x*_tot_ → ∞ will not be given by Eq. (S 45). Since in both cases, for positive and negative *α*, Eq. (S 45) is an approximation for the avalanche part of the lineage size probability density, we call it 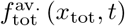.

### B. Most-likely paths

We now derive the most likely path *s_τ_* (*t*) that a critical S-cell population (i.e. *α* = 0) takes from *s*_0_ cells at *t* = 0 to extinction at *t* = *τ*. Since we consider only S-cells and only critical populations, equation 13 reduces a process described by the stochastic differential equation (SDE)

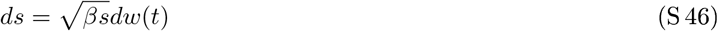

where *w*(*t*) is the standard Brownian motion. If the set of possible paths was finite-dimensional, we could proceed by finding the density *W* defined by equation (S 46) on the set of possible paths starting at *s*_0_ and maximizing *W*(*s_τ_*) subject to *s*_τ_(*τ*) = 0 to find *s_τ_*. It turns out, however, that this approach only cleanly generalizes to infinite-dimensional path spaces in the case of a constant diffusion term [55]. To avoid these technical difficulties, we perform a change of variable to transform equation (S 46) into a process with a constant diffusion term. By setting 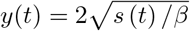 and applying Itô’s lemma we get

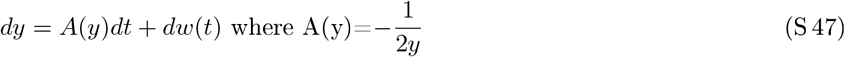

(were once *y* reaches zero, we define it to remain there, despite *A* becoming undefined). For this process, the density functional *W* expressed in terms of its Lagrangian *L* (also called Onsager-Machlup function) is [55]

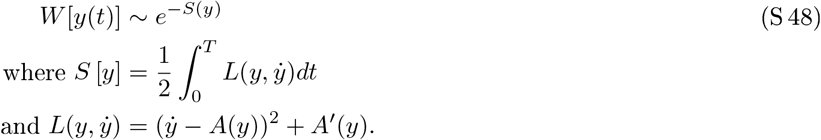

The functional *W* defines a probability density on the set of paths in the following sense: The probability for a random path to lie within a small tube of diameter *ϵ* around a given differentiable path *y* is asymptotically proportional to *W*[*y*]*H*(*ϵ*) for some function *H* independent of *y* (this factorization fails if the diffusion term is not constant [55]). We can therefore proceed as we would in the finite-dimensional case and maximize *W* to find *y_τ_*. Since maximizing W means minimizing the *action* 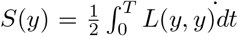, the desired *y_τ_* is found by solving the Euler-Lagrange (EL) equation 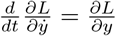. In our case the EL equation yields 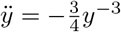 with the general solution

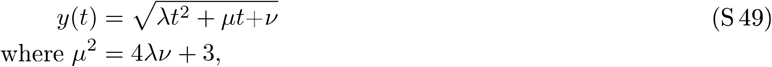

and solving for the boundary conditions 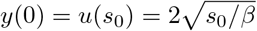 and *y*(*τ*) = 0 yields

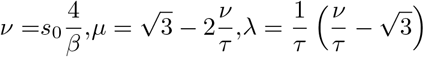

(where we chose the solution that ensures *λt*^2^ + *μt*+ *ν* >= 0 on [0, *τ*]). Inserting these parameter into the general solution (S 49) and transforming back from *y* to *s* produces after some rearrangements the extinction trajectory stated in equation (18).

## Notes

### Competing Interest Statement

The authors have declared no competing interest.

